# Antibacterial Activity of Honey on Wound Isolates from Patients Attending Benue State University Teaching Hospital, Makurdi

**DOI:** 10.1101/2025.09.24.677996

**Authors:** David Tyover Kajo, Martina E Okoh

## Abstract

The medical importance of honey has been documented in many old medical literatures and since ancient times. It has been known to poses antimicrobial properties and wound healing activity; this is mostly due to the enzymatic production of hydrogen peroxide H_2_O_2_, some non-peroxide components and its acidic nature. The current problems with conventional antibacterial agents have led to the choice of honey and other natural products by the populace in treatment of bacterial infections. The present study therefore evaluates the Antibacterial activity of honey on wound isolates from patients attending Benue state university, Makurdi and compared same with ampiclox. Different concentrations (50, 75 and 100%) honey were studied using agar disc diffusion on *Staphylococcus aureus, Escherichia coli, Klebsiella pneumoniae* and *Proteus mirabilis* and data was subjected to chi-square analysis. All bacteria were susceptible to honey at 100% concentration. *Proteus mirabilis* had the highest Z.I (25mm), *S. aureus* (23mm), *E. coli* (22mm) and *Klebsiella* (10mm). Inhibition zone of neat honey and ampliclox on test bacteria were *E. coli* (22, 20mm), *S. aureus* (23, 25mm), *Proteus mirabilis* (25, 23mm) and *Klebsiella* (10, 11mm) respectively. While antibacterial activity at different dilutions was significant, Minimum Inhibitory Concentration (MIC) of honey was determined using broth dilution method and data obtained presented *Proteus mirabilis* as most susceptible organism and *Klebsiella* the least. Honey also had an acidic pH of 4.15 and ash content of 0.029g. The neat concentration (100%) had potency similar but not superior to that of ampiclox. Increase in this branch of alternative medicine and consumption of honey is therefore highly recommended.

## 1. Introduction

Honey is a thick sweet liquid made by honey bees gotten from nectar of flowers. Bees create honey during their routine of gathering nectar from plants. The nectar eventually becomes honey, when the nectar gathered is stored in a hive and special enzymes are added. The honey produced is stored in the small waxy cells of the honey comb. It is a popular sweetener, nontoxic, nonirritant and a common house hold product. Honey is composed mainly of sugars (70 – 80%) such as fructose, sucrose, glucose, etc a low level of water, proteins, hydrogen peroxide and gluconic acid (Agbagwa *et al*., 2010). Honey has high nutritive value as it contains several physicochemical and mineral components (Al-wail, 2005). It is rich in both enzymatic antioxidants and non-antioxidants including catalase, ascorbic acid and flavonoids and alkaloids. Apart from possessing these nutritive qualities, honey had been found useful in management of wound which enables healing and prevents the spread of infection to other body tissues. However, all honeys are not chemically equal and new bioactive components are still being discovered (Kwakman, 2011). The antibacterial properties of honey may be due to its acidic nature and hydrogen peroxide produced enzymically.

The use of traditional medicine to treat infection has been practiced for centuries and honey is one of the oldest traditional medicines considered to be important in the treatment of several human ailments. Honey is hygroscopic in nature and hence speeds up growth of healing tissue and sometimes dries it (Molan *et al*., 2010).

Recent researches have shown that honey is effective in clearing wound infections where other treatments have failed and in preventing sepsis in burns. Antimicrobial agents are essentially important in reducing the global burden of infectious disease, however as resistant pathogens develop and spread, the effectiveness of the antibiotic is diminished (Jande, 2016). This type of bacteria resistance to antimicrobial agents posses a very serious threat to public health and for all kinds of antibiotics, the frequencies of resistance are increasing worldwide (Mandal *et al*., 2010, 2009). Efforts are therefore been made to develop antimicrobial agents from local sources for better chemotherapeutic effects.

This research therefore highlight proximate analysis of honey, isolation and identify pathogens associated with wound and seeks to test the potency and antibacterial property of locally sourced honey against some wound infectious bacteria on wound samples of patients attending Benue State University Teaching Hospital Makurdi, Benue State.

### Objectives

- To determine the antibacterial activity of natural honey against some human pathogens as a contribution to the search of alternative medicine.
- To determine the Minimum Inhibitory Concentration (MIC) and Minimum Bacteriocidal concentration of honey on wound bacteria.
- To determine the major physicochemical components of the honey.

## 2. Materials and Methods

### 2.1 Source of Honey

The honey used for this study was obtained from Adikpo, Kwande Local government area, Benue state. The honey was filtered through a sterile mesh cloth to remove the debris.

### 2.2 Collection of Test Organisms

The bacterial isolate were gotten from Benue state university teaching hospital Makurdi. The isolates; *Staphylococcus aureus, Klebsiella pneumoniae, Escherichia coli* and *Proteus mirabilis* were selected due to their medical importance in causing wound infections.

All organisms were taken to the lab in pure cultures and maintained on nutrient agar slant.

### 2.3 Preparation of Test Bacterial Isolates

Fresh plates of the test bacteria were prepared from the isolated culture obtained on agar slants. Colonies of each of the isolates were picked with an inoculating loop and was then diluted with sterile distilled water to turbidity that matches 0.5McFarland standard (10^5^CFU/ml). 1 ml of the standard inoculums of the different bacterial isolates was used in flooding nutrient agar plates in the agar diffusion method of in vitro antimicrobial susceptibility testing.

### 2.4 Honey Dilution

Honey samples were diluted to different concentrations of 50%, 75% with corresponding volumes of sterile distilled water and undiluted honey (100%) served as neat as described by Yahaya *et al*., (2015).

### 2.5 Antibacterial Assay Using Agar Disc Diffusion Method

Filter paper discs of 6 mm diameter were prepared as described by Mahendran and Kumarasamy, (2015). The discs were impregnated with the different concentrations of each honey 50%, 75%, 100% and ampiclox as control. A sterile cotton swab was dipped into the standardized bacterial suspension and used to evenly inoculate the agar plates. They were allowed to dry for 3 to 5 minutes. Thereafter, all discs were placed on the plates and pressed gently to ensure complete contact with agar. The plates were allowed on the bench for 40 minutes for pre-diffusion, and then incubated at 37^0^C overnight. The resulting zones of inhibition were measured in millimeters.

### 2.6 Determination of Minimum Inhibitory Concentration (Mic) Using Broth Dilution

The minimum inhibitory concentration gave the lowest concentration (highest dilution) of the honey that can inhibit the growth of the test bacteria. This was determined using the broth dilution method as described by Ceyhan and Ugar (2001).

Freshly prepared broth was used in sterile tubes. 1 ml of nutrient broth was put into test tubes number two (2) to test tube number twelve (12). 1 ml of the honey was added to tubes 1 and 2. The honey in tube 2 was therefore diluted 1:2. It was properly mixed then 1 ml was transferred to tube 3 giving 1:4 dilutions. This was continued until the 11^th^ tube from which 1 ml was discarded. The tube 12 which contain only nutrient broth was added to all tubes.

The entire procedure was repeated for all test organisms that were susceptible to honey. The tubes were thoroughly mixed and incubated at 37^0^C for 24 hours, thereafter were usually observed for turbidity after incubation by comparing with control tube.

### 2.7 Determination of Minimum Bacteriocidal Concentration (Mbc) Using Broth Dilution

The MBC is the lowest concentration of a specific antimicrobial agent that kills 99.9% of a given strain of bacteria (Nester *et al*., 2004). This was determined by sub-culturing the tubes from MIC that shows no growth on a fresh honey free medium and incubated for 24 hours. The concentration at which no growth occurs served as the Minimum Bactericidal concentration.

### 2.8 Physicochemical Analysis of Honey

**ph:** the pH of the honey sample was determined using a pH meter

**Ash content:** 10 g of honey sample was weighed into a dried crucible and incinerated in a furnace at 600^0^C to a constant weight.

Ash, % by mass = W2 – W1/M

W2 = weight of Ash + crucible

W1 = weight of Crucible

M = weight of Initial sample before burning

## 3. Results

The data obtained shows inhibitory effect of honey at 50, 75 and 100% concentrations on all the test organisms. The neat Honey (100%) showed the highest antibacterial activity with 25 mm zone of inhibition, while 50% concentration showed the least antibacterial activity with 0 mm inhibition. The result was statistically significant at P<0.05.

The result in Table 2 showed that ampiclox had the highest zone of inhibition against *Staphylococcus aureus* (25mm) when compared with neat honey which had the least against *Klebsiella pneumonia* (10 mm). The neat concentration had potency similar but not superior to that of ampiclox. The result is statistically not significant at P>0.05.

**Table 1:** Antibacterial activity of different concentrations of honey against bacterial isolates.

| Concentration of honey (mg/ml) | *Staphylococcus aureus* (mm) | *Escherichia coli* (mm) | *Klebsiella pneumoniae* (mm) | *Proteus mirabilis* (mm) | Total (mm) |
| --- | --- | --- | --- | --- | --- |
| 100 | 23 | 22 | 10 | 25 | 80 |
| 75 | 19 | 25 | 0 | 21 | 65 |
| 50 | 15 | 0 | 0 | 19 | 34 |
| Total | 57 | 47 | 10 | 65 | 179 |

$\chi^2$ cal = 31.463; p-value= 0.00, p<0.05

**Table 2:**
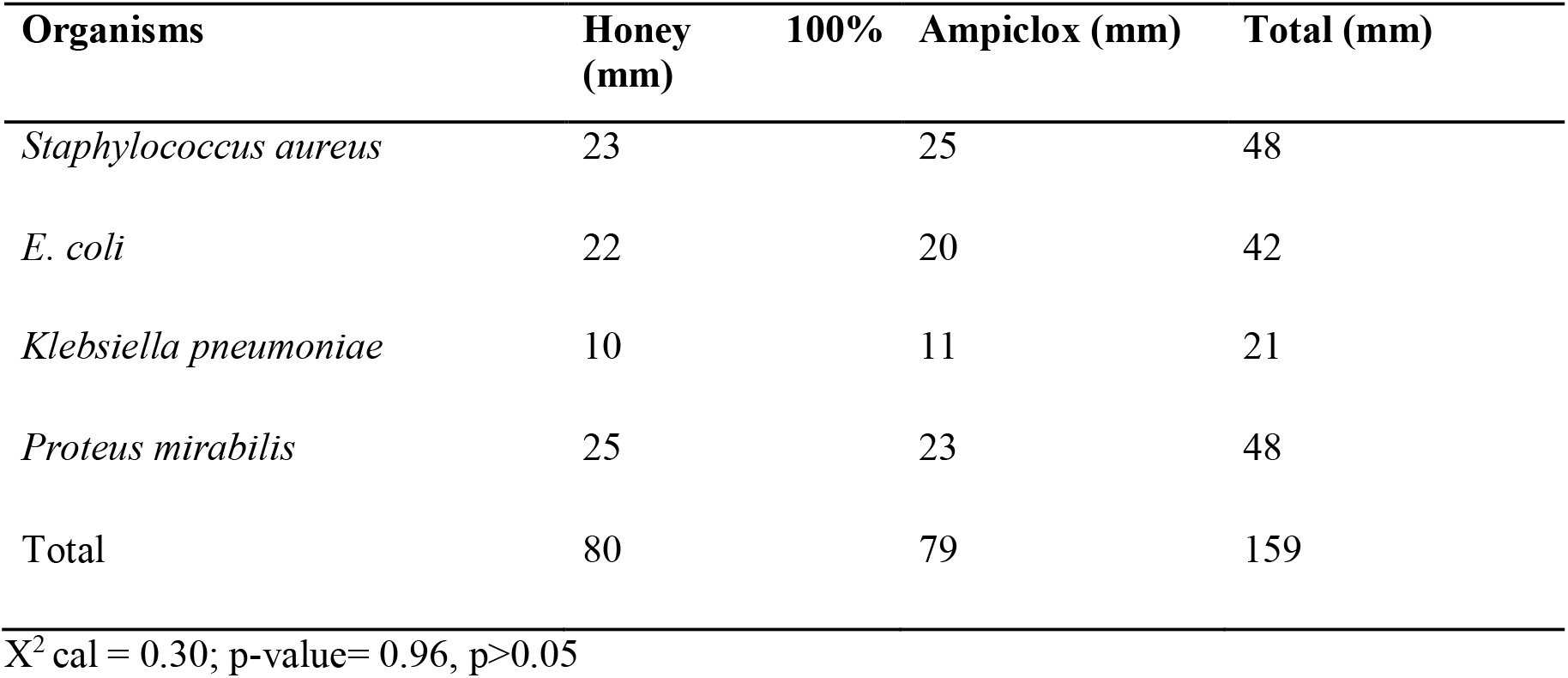
Antibacterial activity of neat against bacterial isolates compared with Ampiclox.

Figure 1 shows the Minimum Inhibitory Concentration (MIC) and Minimum Bacteriocidal Concentration (MBC) of honey. The MIC of honey was found to be more effective on *Proteus mirabilis* (1.56 mg/ml) and least effective on *Klebsiella pneumoniae* (12.5 mg/ml) and *Staphylococcus aureus* (12.5 mg/ml).

**Figure 1:**
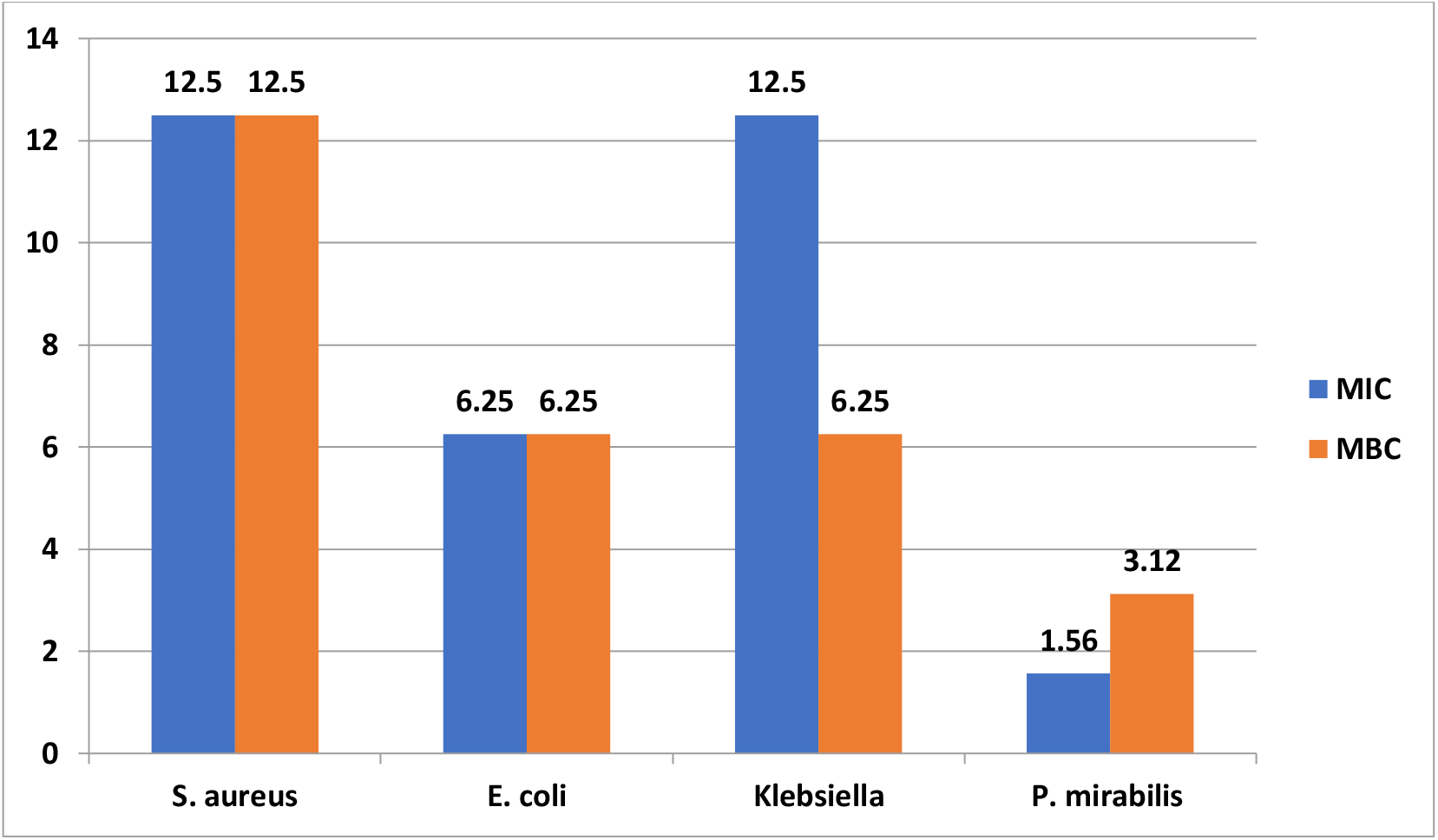
Antibacterial activity of honey using broth dilution method. MIC = Minimum Inhibitory Concentration (mg/ml) MBC = Minimum bacterial Concentration (mg/ml)

Physicochemical analysis of pure honey showed that, honey has a pH of 4.15 and the ash content of 0.029g.

## 4. Discussion

The result obtained from this study shows that the antibacterial activity of undiluted honey was effective against wound pathogenic bacteria which are *Escherichia coli* (22mm), *staphylococcus aureus* (23mm), *Proteus mirabilis* (25mm) and *Klebsiella pneumoniae* (10mm). This agrees with Oyeleke *et al*., 2010 which showed that undiluted honey was also able to inhibit the growth of *E. coli, Staphylococcus aureus, Proteus mirabilis* and *Klebsiella pneumoniae*.

75% concentration of honey also showed antibacterial effect against *Escherichia coli* (25mm), *Staph* (19mm), *Proteus mirabilis* (21mm), *Klebsiella* (0). At 50% concentration, the activity was weak against the pathogens tested. *E. coli* (0), *Staph* (15mm), *Proteus* (19mm), *Klebsiella* (0). This agrees with the findings of Tonks *et al*., 2009 that if honey is diluted, the pH will be so low and the acidity of the honey may not be an effective inhibitor of many species of bacteria. This may probably be responsible for the resistance of *E. coli* and *Klebsiella*. It also agrees with the result of Yahaya *et al*., 2012 which showed that honey can stop growth of bacteria at 50% dilution.

The zone diameter of inhibition (ZDI) obtained in this study for all test bacteria was similar to those obtained by Agbagwa and Frank, 2010 who tested Nigerian honeys from the 4 regions viz; North, West, East and south and found the average ZDI for *S. aureus, P. aeruginosa, E. coli* and *P. mirabilis* to be (5.3-15.6)mm, (14-15.4)mm, (4.4-13.4)mm and (9.1-17)mm respectively and with honey concentrations 80-100%. It again agreed with Agbagwa and Frank, 2010 that the antibacterial effect of honey samples on microorganisms increased as honey concentrations increase.

The MIC of honey on *E. coli* was found to be 6.25mg/ml while that of *S. aureus* and *Klebsiella* was 12.5mg/ml, *Proteus mirabilis* had MIC of 1.56mg/ml. This corroborated with the findings of Oyeleke *et al*., 2010 which reveals that honey can be used as a therapeutic drug against tested bacterial pathogens. Data from MIC suggests that the most susceptible microbe to honey was *Proteus mirabilis* and the order of susceptibility is *Proteus mirabilis* > *S. aureus* > *E. coli* > *Klebsiella pneumoniae*. This was found to be lower than those obtained by Agbaje *et al*., 2006 who recorded MICs of (3.20, 12.80 and 64.0)mg/ml for *Proteus mirabilis, E. coli* and *S. aureus* respectively using honey from Ogun state Nigeria and equal to the ones obtained by Raied, 2009 (with MICs of 12.5 and 1.5mg/ml for *S. aureus* and *Proteus* respectively using local honey from Basrah region, Iraf).

Physicochemical analysis showed that the pH of honey was 4.15, which is low enough to inhibit the growth of pathogens. The optimum pH for growth of bacterial species normally falls between 7.2 and 7.4. The minimum pH values for growth of some common wound infected species as reported by Oyeleke *et al*., 2010 are; E. coli 4.3, *Pseudomonas aeruginosa* 4.4, *Salmonella spp*. 4.0. This pH values signifies that undiluted honey has significant antibacterial factors.

## 5. Conclusion

The crude honey used showed excellent antibacterial activity against *E. coli, S. aureus, Proteus mirabilis* and *Klebsiella pneumoniae* related respectively to urinary tract infection, skin lesion, enteric fever and diarrhea among human patients and thus the honey may be considered against such common infections. Microbial resistance to honey has never been reported which also makes it a very promising antimicrobial against infections of antibiotic-resistant bacteria. The low pH of honey showed that honey possess some acidic quality which provide antibacterial activities against the tested wound pathogens.

## Recommendations

- Further studies which include pharmacological standardization and clinical evaluation on the effect of honey in order to consider it as a preventive and curative measure to the infections caused by the test bacterial strains are encouraged.
- Comprehensive study should be undertaken to determine the various types of honey as it will serve as a guide in choosing the honey most suitable for particular cases.
- Oral consumption of crude honey is also encouraged due to its excellent antibacterial and immunological properties.

## APPENDIX

**Minimum Inhibitory Concentration (MIC) of Honey on test Organisms**.

| Isolates | Neat | 50 | 25 | 12.5 | 6.25 | 3.12 | 1.56 | 0.78 | 0.39 | 0.19 |
| --- | --- | --- | --- | --- | --- | --- | --- | --- | --- | --- |
| *S. aureus* | - | - | - | - | + | + | + | + | + | + |
| *E. coli* | - | - | - | - | - | + | + | + | + | + |
| *K. pneumoniae* | - | - | - | - | + | + | + | + | + | + |
| *P. mirabilis* | - | - | - | - | - | - | - | + | + | + |

(-) No visible growth (not turbid)
(+) Visible growth (turbid)

**Plate 1:**
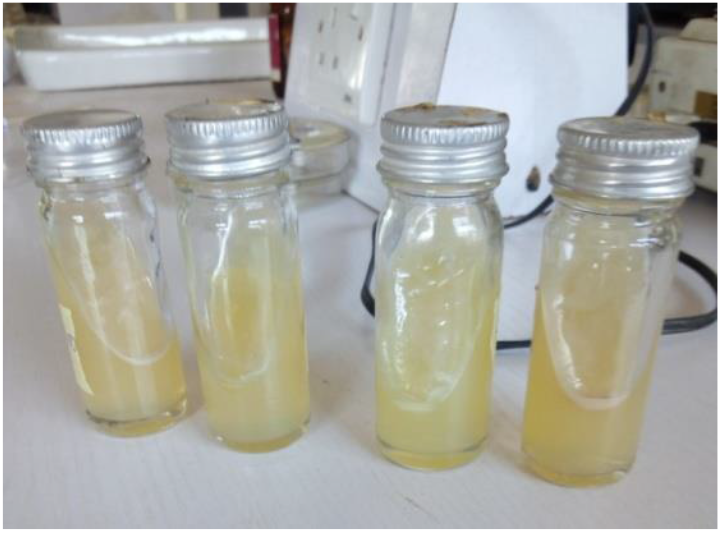
Bacteria isolates on agar slants.

**Plate 2:**
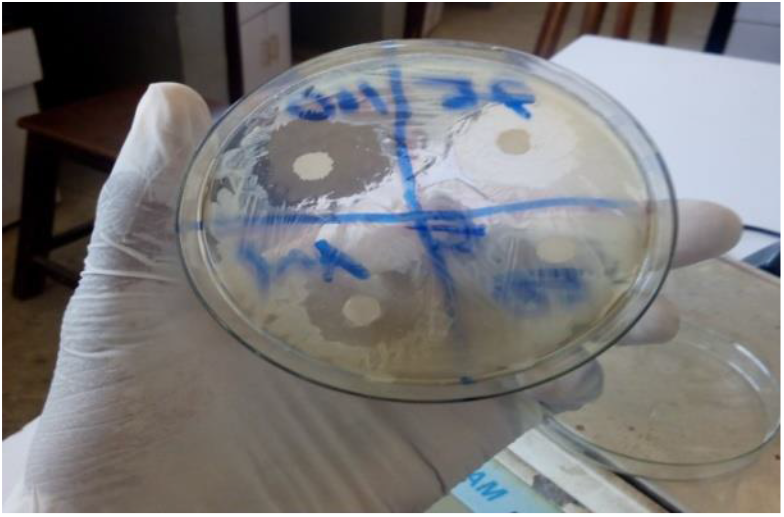
Zone of Inhibition.

**Plate 3:**
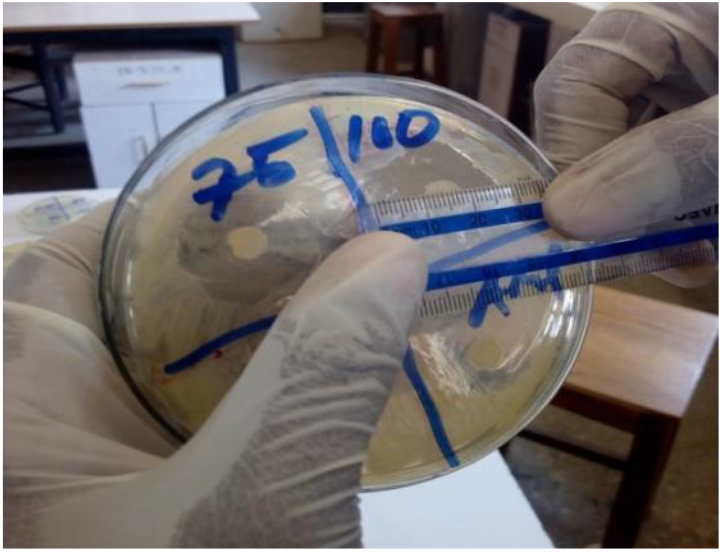
Measurement of Zone of inhibition.

**Plate 4:**
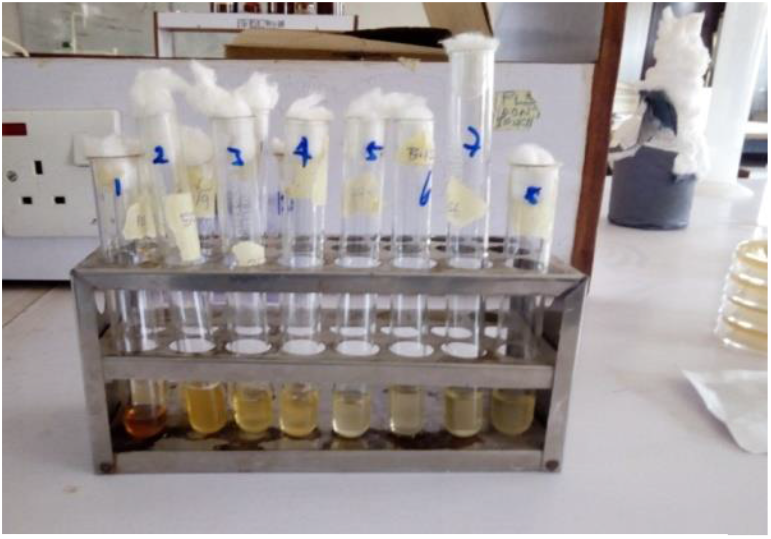
Setup for determination of MIC.

## Notes

### Competing Interest Statement

The authors have declared no competing interest.

